# INeo-Epp: A novel T-cell HLA class-I immunogenicity or neoantigenic epitope prediction method based on sequence related amino acid features

**DOI:** 10.1101/697011

**Authors:** Guangzhi Wang, Huihui Wan, Xingxing Jian, Yuyu Li, Jian Ouyang, Xiaoxiu Tan, Yong Zhao, Yong Lin, Lu Xie

**Affiliations:** College of Food Science and Technology, Shanghai Ocean University, Shanghai, 201306, China; Shanghai Center for Bioinformation Technology, Shanghai Academy of Science and Technology, Shanghai, 201203, China; School of Medical Instrument and Food Engineering, University of Shanghai for Science and Technology, Shanghai, 200093, China; Key Laboratory of Carcinogenesis and Cancer Invasion, Ministry of Education; Key Laboratory of Carcinogenesis, National Health and Family Planning Commission, Xiangya Hospital, Central South University, Changsha,410008, China

## Abstract

In silico T-cell epitope prediction plays an important role in immunization experimental design and vaccine preparation. Currently, most epitope prediction research focuses on peptide processing and presentation, e.g. proteasomal cleavage, transporter associated with antigen processing (TAP) and major histocompatibility complex (MHC) combination. To date, however, the mechanism for immunogenicity of epitopes remains unclear. It is generally agreed upon that T-cell immunogenicity may be influenced by the foreignness, accessibility, molecular weight, molecular structure, molecular conformation, chemical properties and physical properties of target peptides to different degrees. In this work, we tried to combine these factors. Firstly, we collected significant experimental HLA-I T-cell immunogenic peptide data, as well as the potential immunogenic amino acid properties. Several characteristics were extracted, including amino acid physicochemical property of epitope sequence, peptide entropy, eluted ligand likelihood percentile rank (EL rank(%)) score and frequency score for immunogenic peptide. Subsequently, a random forest classifier for T cell immunogenic HLA-I presenting antigen epitopes and neoantigens was constructed. The classification results for the antigen epitopes outperformed the previous research (the optimal AUC=0.81, external validation data set AUC=0.77). As mutational epitopes generated by the coding region contain only the alterations of one or two amino acids, we assume that these characteristics might also be applied to the classification of the endogenic mutational neoepitopes also called ‘neoantigens’. Based on mutation information and sequence related amino acid characteristics, a prediction model of neoantigen was established as well (the optimal AUC=0.78). Further, an easy-to-use web-based tool ‘INeo-Epp’ was developed (available at http://www.biostatistics.online/INeo-Epp/neoantigen.php)for the prediction of human immunogenic antigen epitopes and neoantigen epitopes.

## Introduction

An antigen consists of several epitopes, which can be recognized either by B- or T-cells and/or molecules of the host immune system. However, usually only a small number of amino acid residues that comprise a specific epitope are necessary to elicit an immune response [1]. The properties of these amino acid residues causing immunogenicity are unknown. HLA-I antigen peptides are processed and presented as follows: a). cytosolic and nuclear proteins are cleaved to short peptides by intracellular proteinases; b). some are selectively transferred to endoplasmic reticulum (ER) by TAP transporter, and subsequently are treated by endoplasmic reticulum aminopeptidase;c). antigen presenting cells (APCs) present peptides containing 8-11 AA (amino acid) residues on HLA class I molecules to CD8+ T cells [2]. Researchers can now simulate antigen processing and presentation by computational methods to predict binding peptide-MHC complexes (p-MHC). Several types of software systems have been developed, including NetChop [3], NetCTL [4], NetMHCpan [5], MHCflurry [6]. However, the binding to MHC molecules of most peptides is predicted, only 10%~15% of those have been shown to be immunogenic [7–10]. For neoantigens the result was approximately 5% (range, 1%-20%) due to central immunotolerance [11, 12]. As a result, the cycle for vaccine development and immunization research is extended. Here, we aim to develop a T-cell HLA class-I immunogenicity prediction method to further identify real epitopes/neoepitopes from p-MHC to shorten this cycle.

Many experimental human epitopes have been collected and summarized in the immune epitope database (IEDB) [13], which makes it feasible to mathematically predict human epitopes. However there still exist two limitations: i) a high level of MHC polymorphism produces a severe challenge for T-cell epitope prediction. ii) there is an extremely unequal distribution of data to compare epitopes and non-epitopes. It is not conducive to analyze the potential deviation existing in TCR recognition owing to the presentation of different HLA peptides. A general analysis of all HLA presented peptides, ignoring the specific pattern of TCR recognition of individual HLA presented peptides, may result in a lower predictive accuracy.

With the advances in HLA research, Sette *et al* [14] classified, for the first time, overlapping peptide binding repertoires into nine major functional HLA supertypes (A1, A2, A3, A24, B7, B27, B44, B58, B62). In 2008, John Sidney *et al* [15] made a further refinement, in which over 80% of the 945 different HLA-A and -B alleles can be assigned to the original nine supertypes. It has not been reported whether peptides presented by different HLA alleles influence TCR recognition. Hence, we collected experimental epitopes according to HLA alleles and assume that epitopes belonging to the same HLA supertypes have similar properties.

Moreover, screening for endogenic mutational neoepitopes is one of the core steps in tumor immunotherapy. In 2017, Ott PA *et al.* [16]and Sahin *et al* [17]. confirmed that peptides and RNA vaccines made up of neoantigens in melanoma can stimulate and proliferate CD8+ and CD4+ T cells. In addition, a recent research suggests that including neoantigen vaccination not only can expand the existing specific T cells, but also induce a wide range of novel T-cell specificity in cancer patients and enhance tumor suppression[18]. Meanwhile, a tumor can be better controlled by the combination therapy of neoantigen vaccine and programmed cell death protein 1 (PD-1)/PD1 ligand 1(PDL-1) therapy [19, 20]. Nevertheless, a considerable number of predicted candidate p-MHC from somatic cell mutations may be false positive, which would fail to stimulate TCR recognition and immune response. This is undoubtedly a challenge for designing vaccines against neoantigens.

In our study, based on HLA-I T-cell peptides collected from experimentally validated antigen epitopes and neoantigen epitopes, we aim to build a novel method to further reduce the range of immunogenic epitopes screening based on predicted p-MHC. Finally, a simple web-based tool, INeo-Epp (immunogenic epitope/neoepitope prediction), was developed for prediction of human antigen and neoantigen epitopes.

## Materials and Methods

The flow chart for ‘INeo-Epp’ prediction is shown as follows. (see Figure 1)

**Figure 1:**
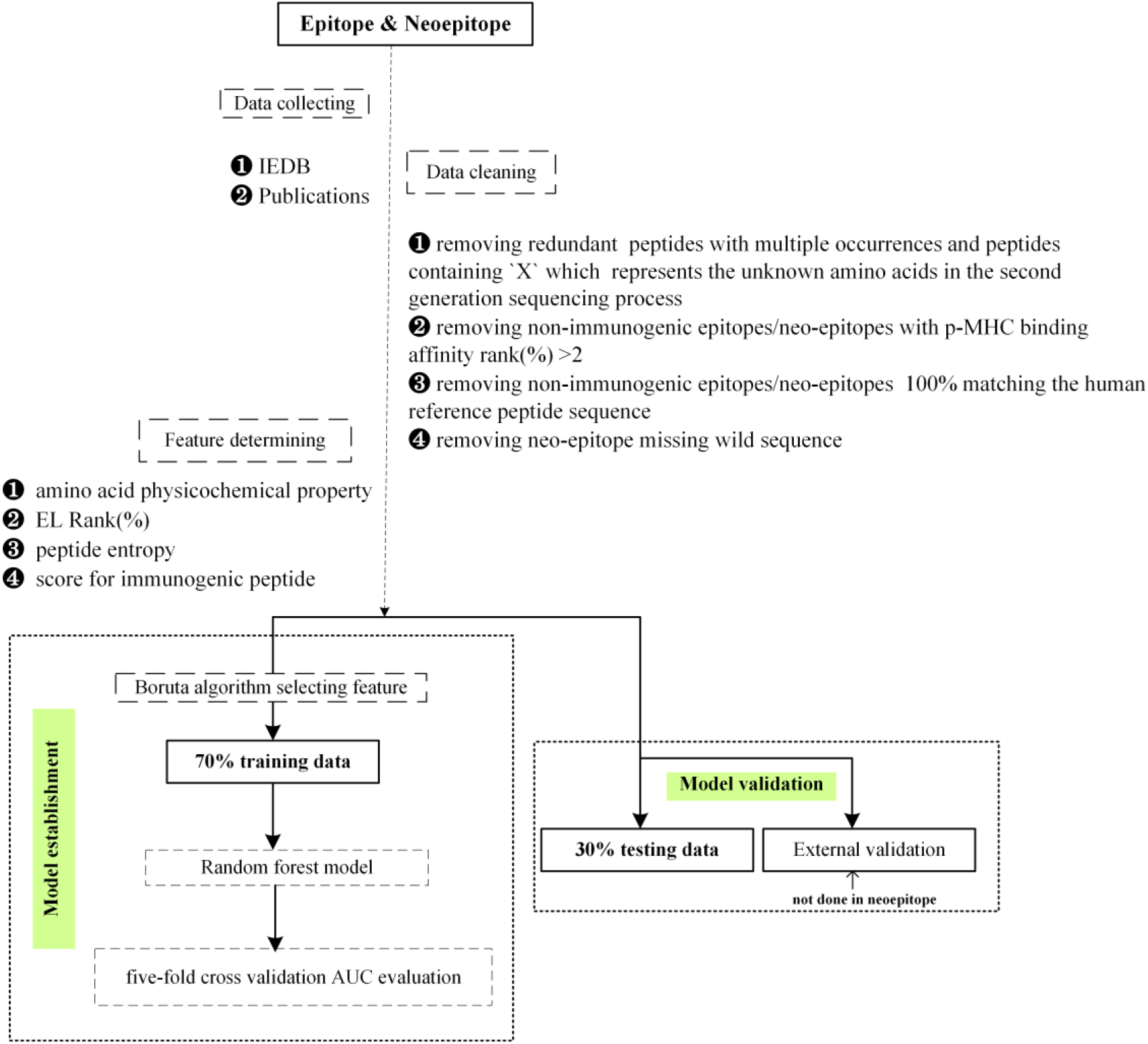
The flow chart for ‘INeo-Epp’ prediction

### Construction of immunogenic and non-immunogenic epitopes

Peptides that can promote cytokine proliferation are considered to be immunogenic epitopes. However, non-immunogenic epitopes may result for the following reasons: a) p-MHC truly unrecognized by TCR; b) peptides not presented by MHC (quantitatively expressed as rank(%)>2, see rank(%) score (below: C24) for details); c) negative selection/clonal presentation induced by excessive similarity to autologous peptides[21]. In this work, to further study the recognition preferences of T cells, peptides with >2 rank(%) were regarded as not in contact with TCR, and sequences 100% matching the human reference peptides (ftp://ftp.ensembl.org/pub/release-97/fasta/homo_sapiens/pep/) were regarded as exhibiting immune tolerance. Hence, we removed these from the definition of non-immunogenic peptides.

### Construction of data sets: epitopes, external validation epitopes and neoepitopes

Antigen epitope data were collected from IEDB (Linear epitope, Human, T cell assays, MHC class I, any disease were chosen). Data collection criteria: each HLA allele quantity >50 and frequency >0.5% (refer to allele frequency database [22]) (Table 1, check Table S1 for detailed information).

**Table 1:**
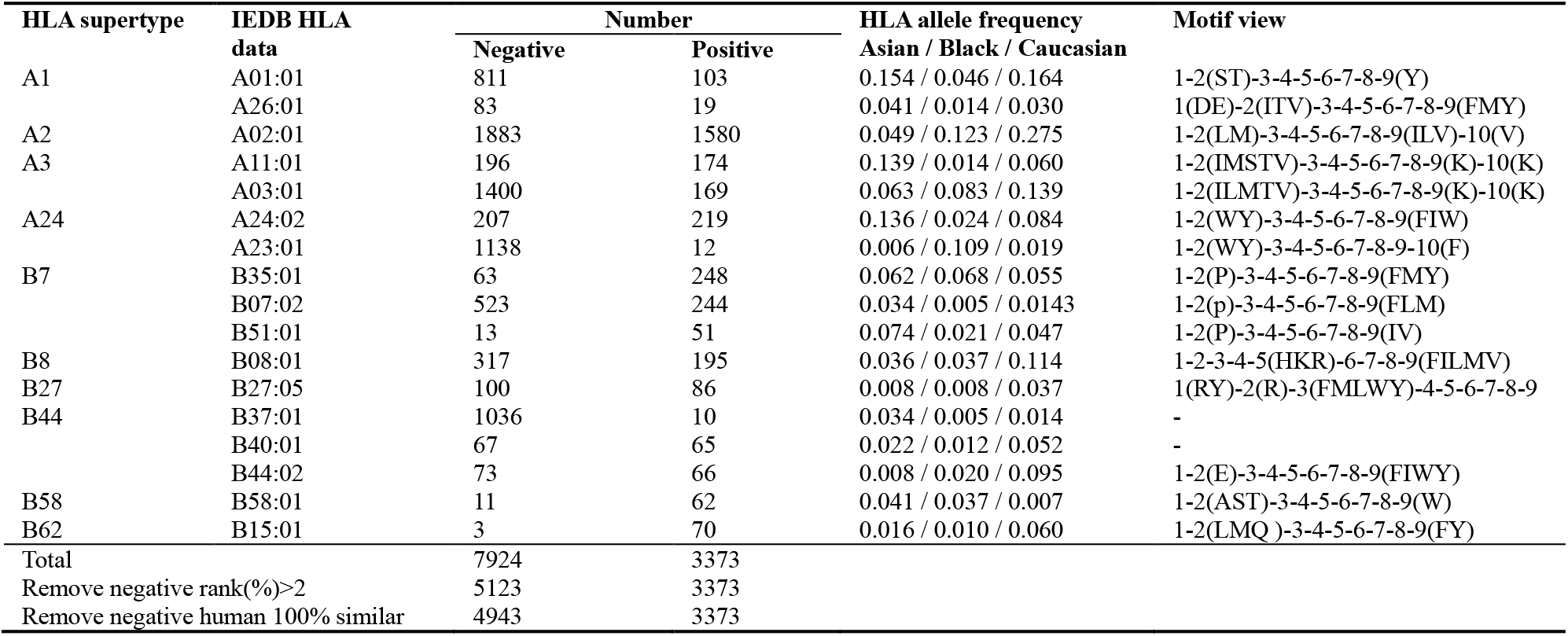
Summary of IEDB epitope data

The external antigen epitope validation set was collected from seven published independent human antigen studies [23–29], consisting of 577 non-immunogenic epitopes and 85 immunogenic epitopes (Table 2, S2 Table)

**Table 2:**
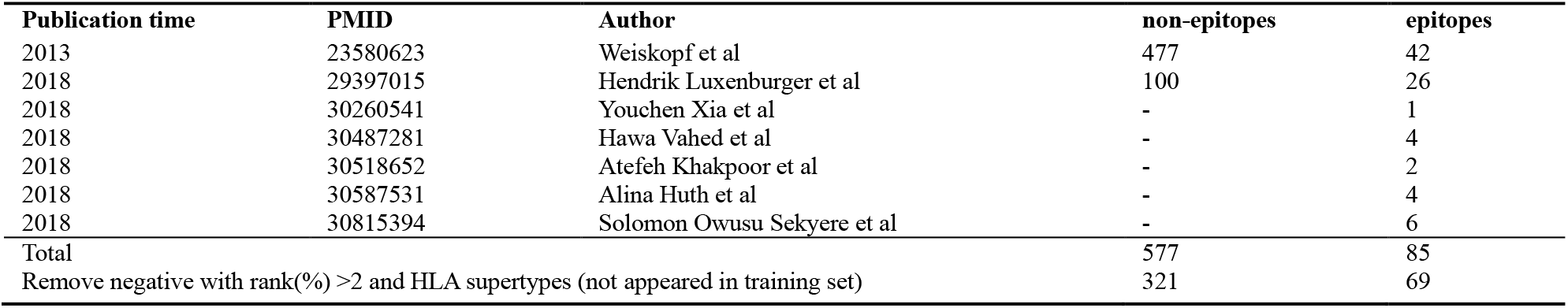
External data included in validation set

Here, we removed peptides for which HLA supertypes do not appear in training set, because we assume peptides belonging to the same HLA supertypes to have similar properties. In the external validation set, some peptides bind to rare HLA supertypes. Their characteristics were not included in the training set. Hence, these peptides in the external validation data might lead to a classification bias.

The neoantigens data were collected from 11 publications [19, 30–39] and IEDB mutational epitopes, and 13 published data sets collected by Anne-Mette B in one publication [40] in 2017 (see Table 3, S3 Table for details) were also included.

**Table 3:**
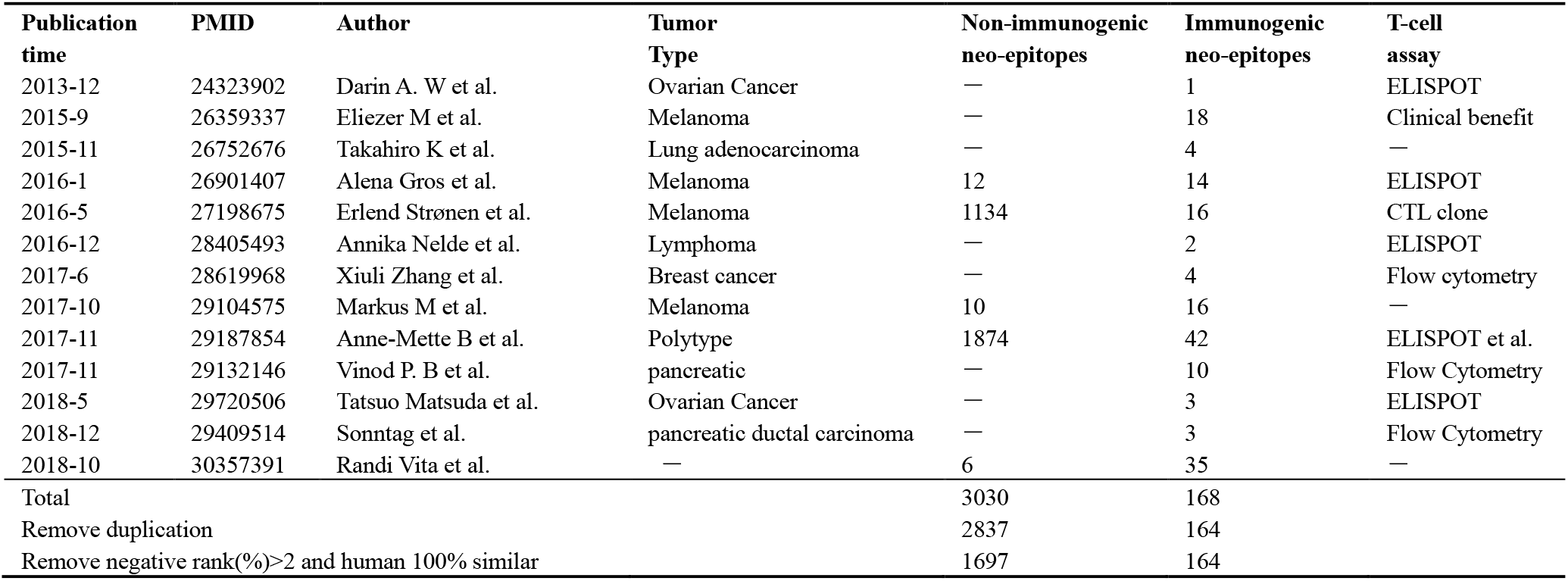
Neoepitopes data included in this study

### Construction of potential immunogenicity feature

#### Characteristics calculation of peptides based on amino acid sequences

The formula for calculating peptide characteristics is shown in (1). P_N_, P_2_, P_C_ (N-terminal, position 2, C-terminal as anchored sites by default) are considered to be embedded in HLA molecules and no contact with TCRs, therefore not evaluated.

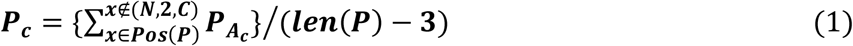

***P***, peptide. ***c***, characteristic. Where ***P_c_*** represents characteristics of peptides. ****A****, amino acid. ****N****, N-terminal in a peptide. ****C****, C-terminal in a peptide. Pos, amino acid position in peptide. Where ***P_Ac_*** represents characteristics of amino acids in peptides.

#### Frequency score for immunogenic peptide (C22)

Amino acid distribution frequency differences between immunogenicity and non-immunogenic peptides at TCR contact sites (excluding anchor sites) were considered as a feature (2).

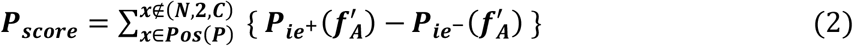

***P*_*ie*_**^+^, immunogenic peptides. ***P*_*ie*_^−^**, non-immunogenic peptides. ***f′*_*A*_**, amino acid frequency in TCR contact position. Where ***P*_*ie*+_ *(f*′_*A*_)** represents frequency of amino acids in immunogenic peptides at TCR contact sites.

#### Calculating peptide entropy (C23)

Peptide entropy [41] was used as a feature (3).

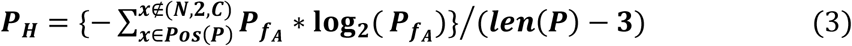

****P_H_,**** peptide entropy. ****f_A_,**** amino acid frequency in human reference peptide sequence. Where ***P_fA_***, represents the frequency in human reference peptide sequence of amino acids in epitope peptides.

#### Rank(%) score (C24)

HLA binding prediction were performed using netMHCpan4.0. rank(%) provides a robust filter for the identification of MHC-binding peptides, in which rank(%) was recommended as an evaluation standard, rank(%)<0.5 as strong binders, 0.5<rank(%)<2 as weak binders, rank(%)>2 as no binders.

### Five-fold cross-validation, feature selection, random forests and ROC generation

The 5-fold cross-validation was implemented in R using the package caret [42] (method = “repeatedcv”, number = 5, repeats = 3). The feature screening results were generated in R using the package Boruta [43] (a novel random forest based feature selection algorithm for finding all relevant variables, which provides unbiased and stable selection of important and non-important attributes from an information system. It iteratively removes the features which are proven by a statistical test to be less relevant than random probes. It uses Z score (computed by dividing the average loss by its standard deviation) as the importance measure and it takes into account the fluctuations of the mean accuracy loss among trees in the forest). R package randomForest [44] was used for training data (the R language machine learning package caret provides automatic iteration selection of optimal parameters, mtry=15 for antigen epitope, mtry=14 for neoantigen epitope, the remaining parameters use default values). R package ROCR [45] was used for drawing ROC.

### Web tool implementation

The front-end of Ineo-Epp was constructed via HTML/JavaScript/CSS. The back end was written in PHP, connecting the web interface and Apache web server. A python script was used for calculating peptide characteristics and extracting mutation information. Models were built using R.

## Results

Ultimately, 11,297 validated epitopes and non-epitopes with the length of 8-11 amino acids were collected from IEDB. T-cell responses included activation, cytotoxicity, proliferation, IFN-γ release, TNF release, granzyme B release, IL-2 release, IL-10 release, etc. Seventeen different HLA alleles were collected (Fig 2A), and the detailed antigen length distribution is shown in (Fig 2B). Additionally, we collected the neoantigen data from 12 publications, including 2837 non-neoepitopes and 164 neoepitopes (Fig 2C), and the detailed neoantigen length distribution is shown in (Fig 2D).

**Figure 2:**
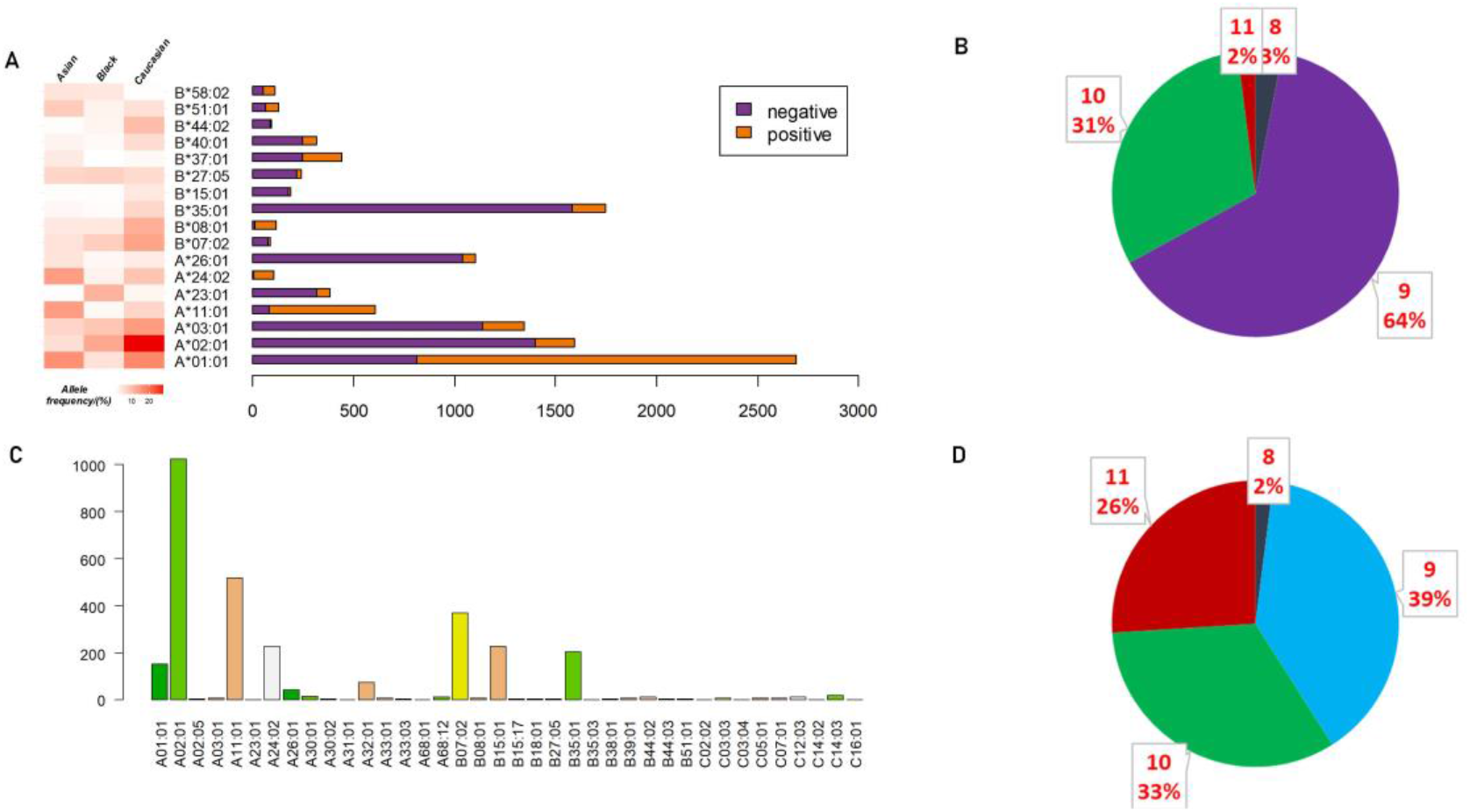
Epitope/neoepitope peptides composition and amino acid lengths distribution. (a) Detailed data distribution of seventeen HLA alleles of antigen peptides and proportion of each HLA allele (positive and negative) epitopes and the corresponding HLA frequency in Asian, Black, Caucasian. (b) Proportion of antigen peptides of 8-11 AA lengths. (c) Data distribution of HLA alleles of neoantigen peptides. (d) Proportion of neoantigen peptides of 8-11 AA lengths.

The TCR contact position plays a crucial role in the analysis of immunogenicity, as TCRs might be more sensitive to some amino acids, the amino acids preference in antigen epitope peptide and antigen non-epitope peptide was further analyzed after excluding anchor sites (N-terminal, position 2, C-terminal) (Fig 3). We found that TCRs tend to identify hydrophobic amino acids. For example, 3/4 hydrophobic amino acids (L, W, P, A, V, M) occur more frequently in immunogenicity epitopes. Charged amino acids (*e.g*. D, K) are enriched in non-epitopes whereas the rest of charged amino acids (R, H, E) show no difference. Based on the result in figure 3, the amino acid distribution difference at the TCR contact sites was regarded by us as one of the immunogenicity features (*i.e.* Frequency score for immunogenic peptide (C22)).

**Figure 3:**
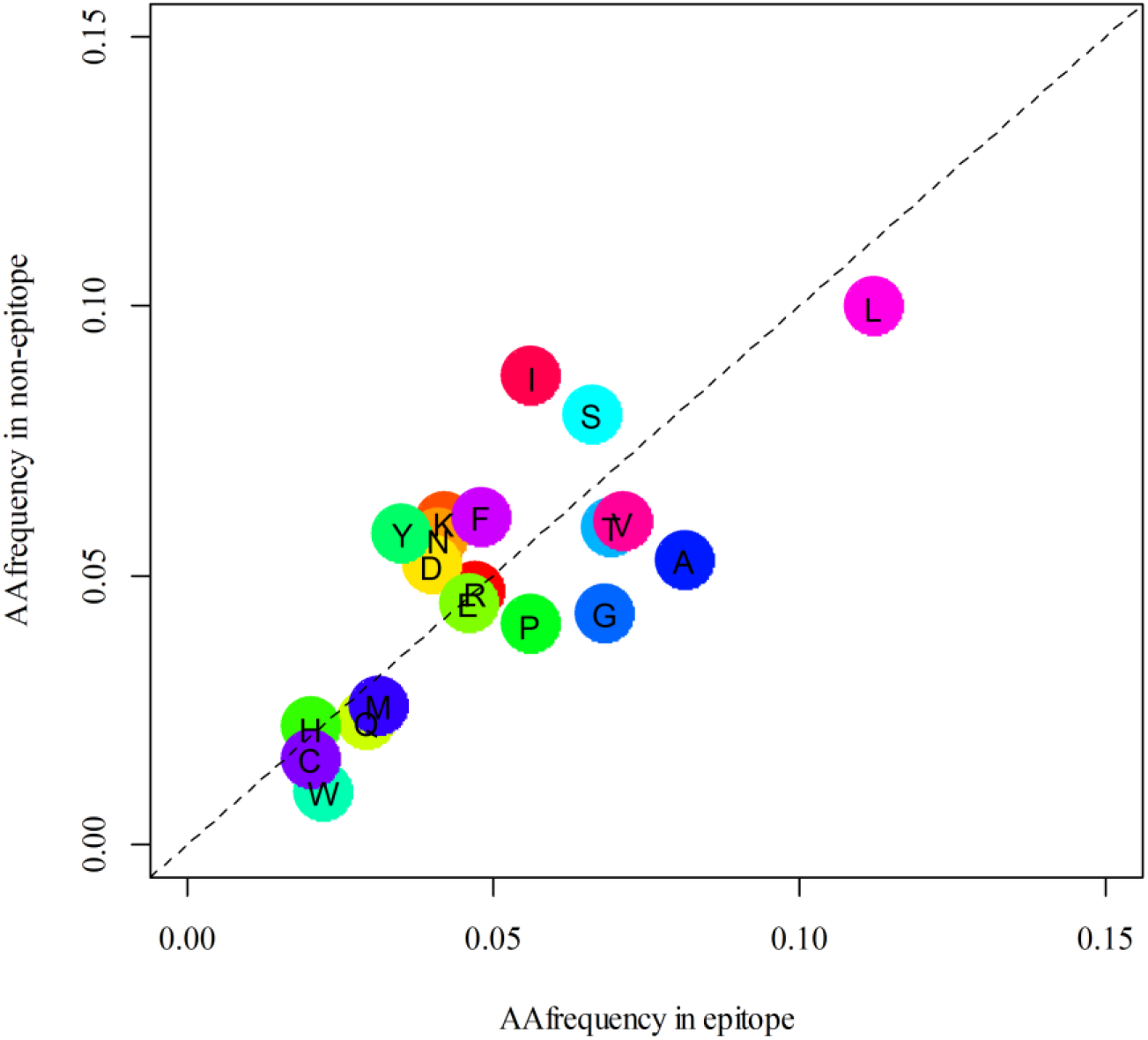
Antigen epitope amino acid distribution frequency in TCR contact site of epitopes and non-epitopes. Frequency distribution of amino acids at TCR contact sites in antigen epitope and non-epitope peptides, and the amino acids below the dotted line are preferred by the epitope.

### Classification prediction model for antigen epitopes

We constructed the features of peptides on the basis of the characteristics of amino acids (see Materials and Methods section: Characteristics Calculation of peptides based on amino acids). All amino acid characteristics were selected from Protscale [46] in ExPASy (SIB bioinformatics resource portal). The 21 involved features are as follows: Kyte–Doolittle numeric hydrophobicity scale (C1) [47], molecular weight (C2), bulkiness (C3) [48], polarity (C4) [49], recognition factors (C5) [50], hydrophobicity (C6) [51], retention coefficient in HPLC (C7) [52], ratio hetero end/side (C8)[49], average flexibility (C9) [53], beta-sheet (C10) [54], alpha-helix (C11) [55], beta-turn (C12) [55], relative mutability (C13) [56], number of codon(s) (C14), refractivity (C15) [57], transmembrane tendency (C16) [58], accessible residues (%) (C17) [59], average area buried (C18) [60], conformational parameter for coil (C19) [55], total beta-strand (C20) [60], parallel beta-strand (C21) [61] (see Table S4 in detail). Also, frequency score for immunogenic peptide (C22), peptide entropy (C23) and rank(%) (C24) were also taken into consideration. Together, 24 immunogenic features were collected, and all features were retained for antigen epitopes prediction after screening using the R package Boruta. Compared with other characteristics, the frequency score for immunogenic peptide and rank(%) have higher impact, suggesting they have more significant influence on antigen epitopes classification (Figure 4A).

**Figure 4:**
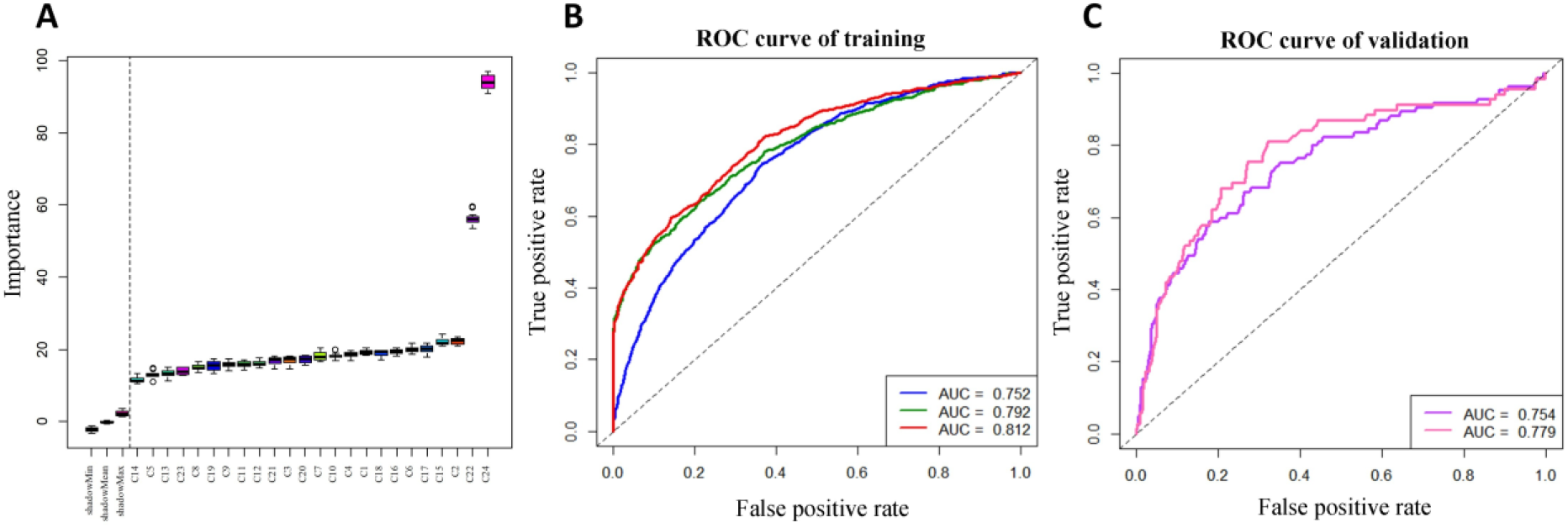
Feature selection in antigen epitopes and ROC curves of antigen epitopes classification. (a)Peptide features: Twenty four features were screened and we defined the features on the right of the dotted line as being effective. (b)Trained model: The line in blue represents antigen epitopes without screening; the line in green represents selection with the deletion of rank(%)>2 non-epitope; and the line in red represents selection with the deletion of the non-epitopes 100% matching human reference peptide sequence. (c)External validation: The ROC curves for the external verification set, line in purple represents modeling using antigen epitopes without filtering, the line in pink represents modeling using antigen epitopes removing non-epitopes which rank(%)>2 and HLA for which supertypes not appearing in training set.

The receiver operator characteristic (ROC) curve of models are shown in Fig 4. The five-fold cross validation AUC was 0.81 in the prediction model for antigen epitope (line in red Fig 4B) and the externally validated (see table 2) AUC was 0.75 (line in purple Fig 4C). Here, we tried to remove peptides for which HLA supertypes not appearing in training set from the externally validated antigen data and, the AUC, specificity, and sensitivity were increased to 0.78, 0.71, and 0.72, respectively. (line in pink Fig4 C). This, to some extent, verifies our conjecture about TCR specific recognition of different HLA alleles presenting peptides.

### Classification prediction model for neoantigen epitopes

Neoantigens derived from somatic mutations are different from the wild peptide sequences. Therefore, some mutation-related characteristics were also taken into account. For instance, difference in hydrophobility before and after mutation (C25), differential agretopicity index (DAI, C26) [62] and whether the mutation position was anchored (C27). Finally, 27 features were selected for the neoantigen epitope prediction model. However, only 25 neoantigen related features were retained after running Boruta, because C25 and C27 were removed. Also, rank(%) showed a marked effect (Fig 5A). in the five-fold cross-validation of the prediction model for neoantigen epitopes, AUC was 0.78 (Fig 5B).

**Figure 5:**
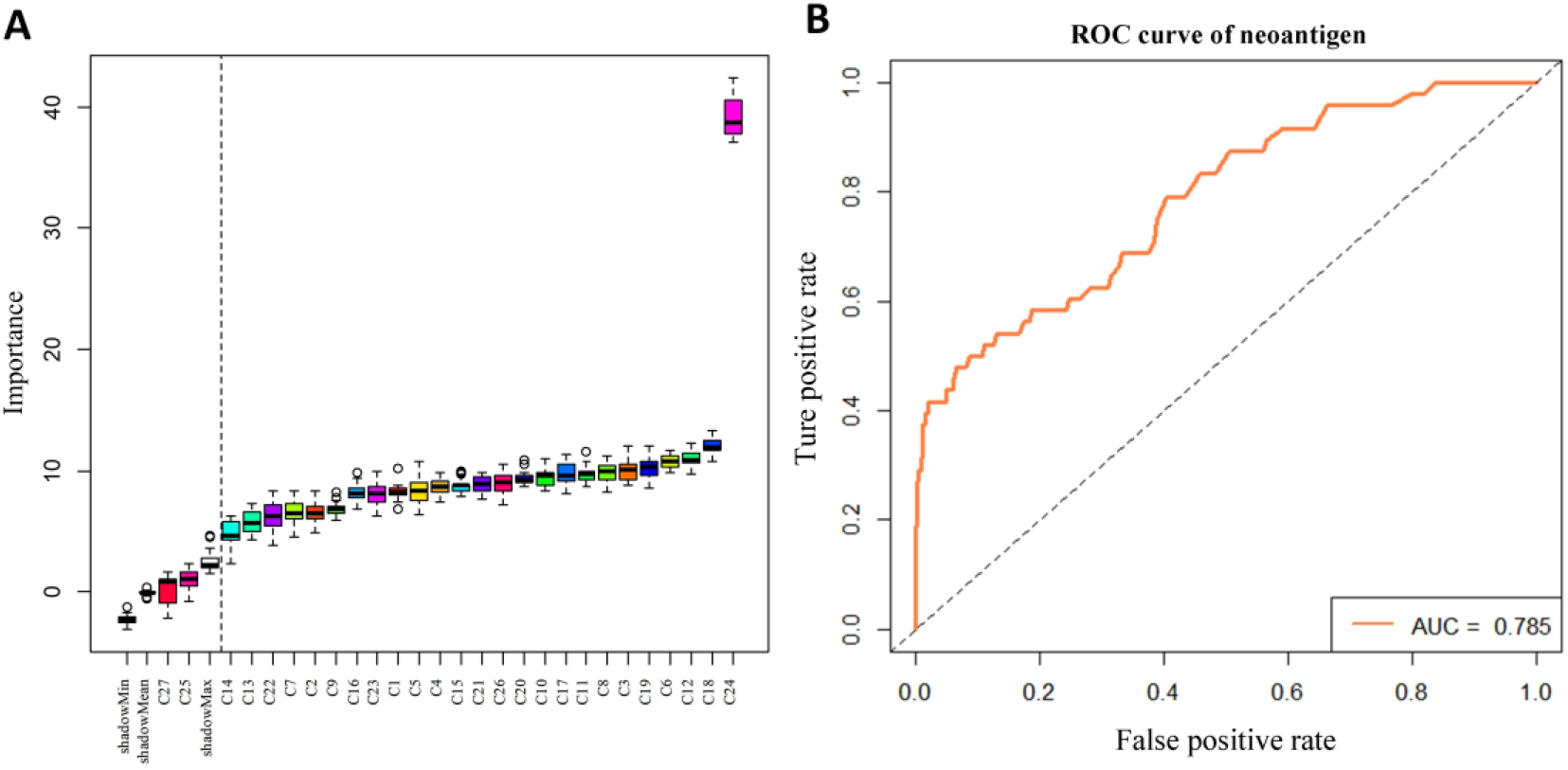
Feature selection in neoantigen epitopes and ROC curves of neoantigen epitopes classification. (a) Twenty seven features were screened and the 25 features on the right of the dotted line were reserved for modeling using a random forest algorithm. (b) ROC curves of neoantigen epitopes classification.

### Web server for TCR epitope prediction

Based on these above-mentioned validated features, we established a web server for TCR epitope prediction, named ‘INeo-Epp’. This tool can be used to predict both immunogenic antigen and neoantigen epitopes. For antigen, the nine main HLA supertypes can be used. We recommend the peptides with the lengths of 8-12 residues, but not less than 8. N-terminal, position 2, C-terminal were treated as anchored sites by default. A predictive score value greater than 0.5 is considered as immunogenicity (Positive-High),the score between 0.4-0.5 is considered as (Positive-Low),the score less than 0.4 is considered as (Negative-High).It is critical to make sure that HLA-subtype must match your peptides(rank(%)<2). Where HLA-subtypes mismatch, the large deviation of rank(%) value may strongly influence the results. Additionally, the neoantigen model requires providing wild type and mutated sequences at the same time to extract mutation associated characteristics, and currently only immunogenicity prediction for neoantigens of single amino acid mutations are supported. Users can choose example options to test the INeo-Epp (http://www.biostatistics.online/INeo-Epp/neoantigen.php).

## Discussion

Due to the complexity of antigen presenting and TCR binding, the mechanism of TCR recognition has not been clearly revealed. In 2013, J. A. Calis [63] developed a tool for epitope identification for mice and humans (AUC = 0.68). Although mice and human beings are highly homologous, the murine epitopes may very likely cause limitations in identifying human epitopes. Inspired by J. A. Calis, our research here focused on human beings’ epitopes and has been conducted in a larger data set.

By analyzing epitope immunogenicity from the perspective of amino acid molecular composition, we observed that TCRs do have a preference for hydrophobic amino acid recognition. For short peptides presented by different HLA supertypes, TCRs may have different identification patterns. The immunogenicity prediction based on all HLA-presenting peptides may affect the accuracy of the prediction results. That is, if the prediction could focus on specified HLA-presenting peptides the results may improve. Therefore in our work we used HLA supertypes to improve the prediction of HLA-presenting epitopes, including antigen epitopes and neoantigen epitopes, for a better recognition by TCRs. At present, neoantigen epitopes that can be collected in accordance with the standard for experimental verification are too few, the data of positive and negative neoantigens are unbalanced, and there is not enough data to be used for external verification set. In the future, we will continue to refine and expand our training and verification datasets. Recently, Céline M. Laumont [64] demonstrated that noncoding regions aberrantly expressing tumor-specific antigens (aeTSAs) may represent ideal targets for cancer immunotherapy. These epitopes can also be studied in the future. Increased epitope data may also help empower the prediction of potentially immunogenic peptides or neopeptides.

## Conclusions

Neoantigen prediction is the most important step at the start of preparation of neoantigen vaccine. Bioinformatics methods can be used to extract tumor mutant peptides and predict neoantigens. Most current strategies aimed at ended in presenting peptides predictions and among the results of these predictions, probably only fewer than 10 neoantigens might be clinically immunogenic and produce effective immune response. It is time-consuming and costly to experimentally eliminate the false positively predicted peptides. Our methods as developed in this study and the INeo-Epp tool may help eliminate false positive antigen/neoantigen peptides, and greatly reduce the amount of candidates to be verified by experiments. We believe that in the age of biological systems data explosion, computational approaches are a good way to enhance research efficiency and direct biological experiments. With the development of machine learning and deep learning, we expect the prediction of epitope immunogenicity will be continually improved.

In summary, this study provides a novel T-cell HLA class-I immunogenicity prediction method from epitopes to neoantigens, and the INeo-Epp can be applied not only to identify putative antigens, but also to identify putative neoantigens.

It needs to be stated here that we published the preprint [65] of this article in July 2019.This is a modified version.

## Data Availability

The data used to support the findings of this study are included within the supplementary information file(s).

## Acknowledgments

We sincerely thank Drs. Menghuan Zhang, Hong Li and Qibing Leng for valuable discussion. We also acknowledge Dr. Michael Liebman for his critical reading and editing.

## Supplementary Material

S1 Table **IEDB antigen epitopes summary**. Detailed description of 17 HLA molecules collected from IEDB. (XLSX)

S2 Table **External validation antigen epitopes summary**. Epitope details of 7 publications. (XLSX)

S3 Table **Neoantigen epitopes summary**. Epitope details of 13 publications. (XLSX)

S4 Table **Summary of amino acid characteristics**. For all amino acid characteristics (n=21) that are described in the ExPASy. (XLSX)

